# Cryptic genetic variation underpins rapid adaptation to ocean acidification

**DOI:** 10.1101/700526

**Authors:** M. C. Bitter, L. Kapsenberg, J.-P. Gattuso, C. A. Pfister

## Abstract

Global climate change has intensified the need to assess the capacity for natural populations to adapt to abrupt shifts in the environment. Reductions in seawater pH constitute a conspicuous stressor associated with increasing atmospheric carbon dioxide that is affecting ecosystems throughout the world’s oceans. Here, we quantify the phenotypic and genetic modifications associated with rapid adaptation to reduced seawater pH in the marine mussel, *Mytilus galloprovincialis*. We reared a genetically diverse larval population in ambient and extreme low pH conditions (pH_T_ 8.1 and 7.4) and tracked changes in the larval size and allele frequency distributions through settlement. Additionally, we separated larvae by size to link a fitness-related trait to its underlying genetic background in each treatment. Both phenotypic and genetic data show that *M. galloprovincialis* can evolve in response to a decrease in seawater pH. This process is polygenic and characterized by genotype-environment interactions, suggesting the role of cryptic genetic variation in adaptation to future climate change. Holistically, this work provides insight into the processes underpinning rapid evolution, and demonstrates the importance of maintaining standing variation within natural populations to bolster species’ adaptive capacity as global change progresses.

## Introduction

A fundamental focus of ecological and evolutionary biology is determining if and how natural populations can adapt to rapid changes in the environment. Recent efforts that have combined natural population censuses with genome-wide sequencing techniques have shown that phenotypic changes due to abrupt environmental shifts oftentimes occur concomitantly to signatures of selection at loci throughout the genome^1–4^. These studies demonstrate the importance of standing genetic variation in rapid evolutionary processes^5^, and challenge classical population genetic theory, which assumes that most genetic variation has a small effect on fitness and that selective forces alter this variation gradually over a timescale of millennia^6,7^. Theoretical^8,9^ and experimental studies^10–13^ have further shown that rapid adaptation via standing variation is oftentimes characterized by genotype-environment interactions, in which a particular genetic background is most fit in one environment, while an alternate genetic background leads to a fitness advantage when the environment shifts^14^.

Recently, it has been suggested that genotype-environment interactions during extreme stress or exposure to novel environmental conditions are underpinned by “cryptic genetic variation”, defined as a subclass of standing genetic variation with a conditional effect, such that it becomes adaptive in a new environment^14,15^. The ways in which cryptic genetic variation is maintained and ultimately influences evolutionary dynamics has been explored in theory^9,16^. But, the relative importance of cryptic variation in nature has yet to be robustly demonstrated, particularly within the context of non-model and ecologically important species^14^. Even amidst its suggested ecological importance to colonization of novel habitats^12^, empirical validation of its presence and role is limited, and has almost exclusively focused on prokaryotic systems^10,13,17^ and model species, such as *Drosophila* and *Arabidopsis*^18,19^. Still, existing empirical work has provided initial evidence that cryptic variation may allow populations to circumvent a fitness valley during an evolutionary response, thereby preventing severe population bottlenecks during rapid adaptation^14^. Confirming the role of cryptic genetic variation in rapid adaptation is especially relevant given the threat of global climate change, as natural populations become increasingly exposed to environmental conditions that exceed contemporary variability. The extent to which current levels of genetic variation will facilitate the magnitude and rate of adaptation necessary for species persistence is unclear^20,21^.

In marine systems one pertinent threat is ocean acidification, the global-scale decline in seawater pH driven by oceanic sequestration of anthropogenic carbon dioxide emissions^22^. The current rate of pH decline is unprecedented in the past 55 million years^23^, and lab-based studies have shown negative effects of expected pH conditions on a range of fitness-related traits (e.g., growth, reproduction, and survival) across life-history stages and taxa^24^. Marine bivalves are one of the most vulnerable taxa to ocean acidification^25,26^, particularly during larval development^27^. The ecologically and economically valuable Mediterranean mussel, *Mytilus galloprovincialis*, is an exemplary species for studying the effects of ocean acidification on larval development. Low pH conditions reduce shell size and induce various, likely lethal, forms of abnormal larval development^28,29^. Sensitivity to low pH, however, can vary substantially across larvae from distinct parental crosses, suggesting that standing genetic variation could fuel an adaptive response to ocean acidification^29^.

Here, we explored the potential for, and dynamics of, rapid adaptation to ocean acidification in *M. galloprovincialis*. We tracked the phenotypic distributions and trajectories of 29,400 single nucleotide polymorphisms (SNPs), from the embryo stage through larval pelagic growth and settlement in a genetically diverse larval population, within artificially imposed ambient (pH_T_ 8.05) and extreme low pH treatment (pH_T_ 7.4). To test for a genotype-environment interaction underpinning adaptation, we associated shell size, a trait negatively correlated with fitness-related abnormalities^29^, to its underlying genetic background in each pH treatment. The results presented demonstrate the capacity for natural populations to adapt to rapid environmental change, and suggest that this process will be fueled by cryptic genetic variation.

## Methods Summary

We quantified the effects of low pH exposure on phenotypic and genetic variation throughout development in a single population of *M. galloprovincialis* larvae (Fig. 1). Larvae were reared in ambient and low pH and (i) shell size distributions were quantified on days 3, 6, 7, 14, and 26; (ii) SNP frequencies across the species’ exome were estimated on days 6, 26, and 43; and (iii) the genetic background of shell size was determined in each treatment to assess the presence of a genotype-environment interaction. To generate a starting larval population representative of the standing genetic variation within a wild population of *M. galloprovincialis*, 16 males were crossed to each of 12 females, hereafter referred to as the founding individuals (*N* = 192 unique crosses). The resulting larval population was reared in an ambient (pH_T_ 8.05, *N* = 6 replicate buckets) and low pH treatment (pH_T_ 7.4-7.5, *N* = 6 replicate buckets). While the low pH treatment falls outside the range of annual variability the population currently experiences (pH_T_ ∼7.8-8.1)^29^, and exceeds the −0.4 pH_T_ units expected globally by 2100^22^, normal development of *M. galloprovincialis* larvae can occur at this pH^29^. We thus expected, *a priori*, that this value would effectively reveal the presence of cryptic variation underpinning low pH tolerance.

**Figure 1.**
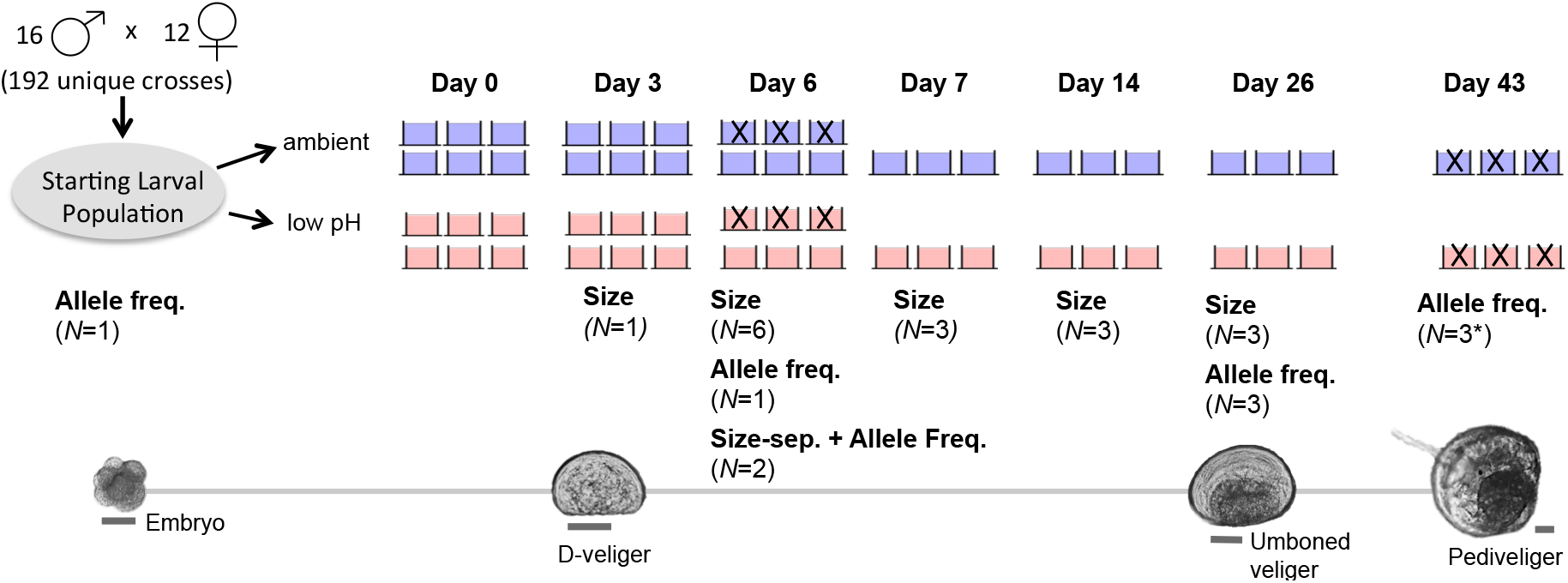
Experimental schematic depicting pictures of larvae at key developmental points, cross design, and replication and sampling strategy throughout the experiment. Scale bar for larval pictures set at 50 μm. Replicate buckets marked with an “X” were destructively sampled (i.e. all larvae removed/preserved) and thus absent from the experimental system on subsequent sampling days. *Allele frequency data from two replicate buckets in the ambient treatment was generated on day 43, as the third replicate bucket to optimize protocol for sampling settled larvae

## Results

### Phenotypic Trajectories

As expected, shell size was significantly affected by pH treatment throughout the experiment (linear mixed effects model: *p* = 0.029), and shell length of low pH larvae was 8% smaller than that of larvae reared in ambient pH on days 3 and 7. Shell length was affected by the interaction of day and treatment (linear mixed effects model: *p* < 0.001), indicating treatment specific growth patterns. From days 7 to 26 the size distributions in each treatment began to converge, with larvae in low pH becoming only 2.5% smaller than those cultured in the ambient treatment by day 26 (Fig. 2).

**Figure 2.**
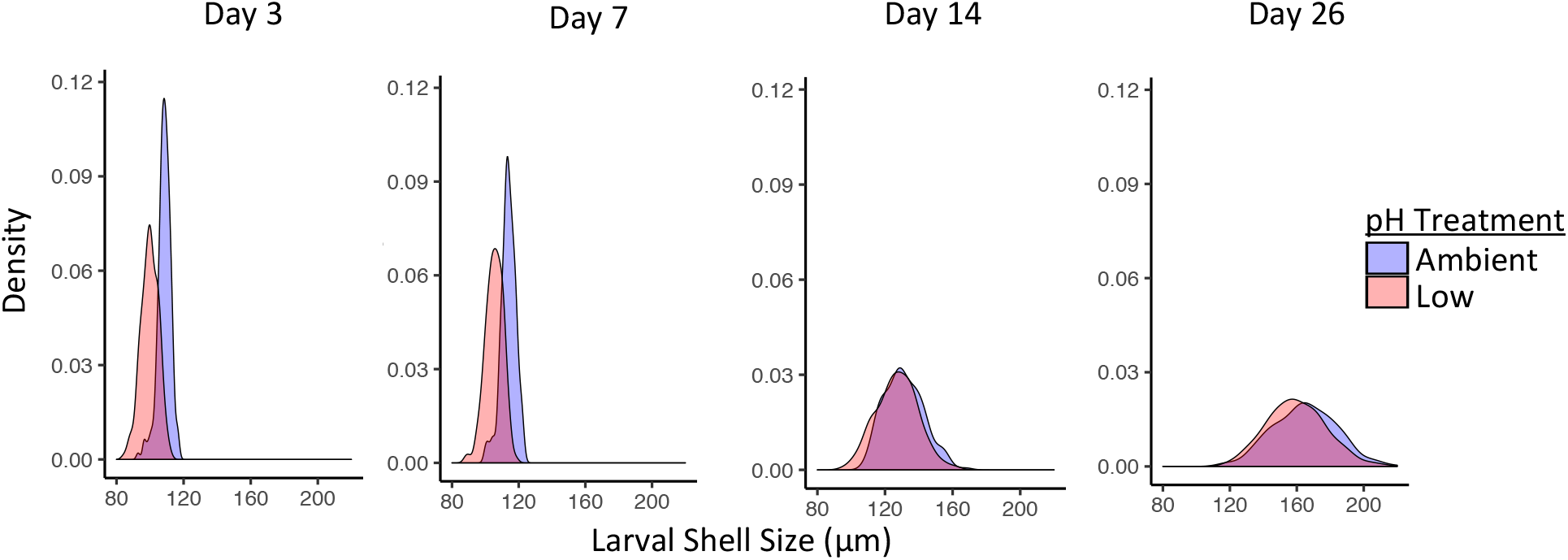
Larval size distributions throughout the shell growing period. Larval size was significantly affected by treatment (p = 0.029) and the interaction of day and treatment (p < 0.001) throughout the shell growing period.

### Changes in Genetic Variation

We identified 29,400 SNPs across the species exome that were present within the larval population across all sampling days and treatments. To link the observed phenotypic trends in each treatment to changes in this variation, we analyzed the SNP’s using principal component analysis (PCA), outlier loci identification, and a statistical metric of genomic differentiation (F_ST_). For both ambient and low pH treatments, all analyses indicated increasing genomic differentiation of the larval cultures away from the day 0 larval population. This trend is visually apparent in the principal component analysis (PCA), which incorporated allele frequency data from all larval samples collected during the pelagic stage and settlement (excluding size-separated groups) (Fig. 3). At later days (e.g. days 26 and 43), there is an observed increase in Euclidian distance among samples. This may be driven, at least in part, by selection-induced declines in larval population survival throughout the pelagic phase, and an associated increase in the influence of allele frequency “drift” among replicate buckets. Observations of increased larval mortality throughout the experiment (indicated via empty D-veliger shells in buckets) corroborated these trends, though we were unable to quantify larval mortality (MCB, *pers. obs.*).

**Figure 3.**
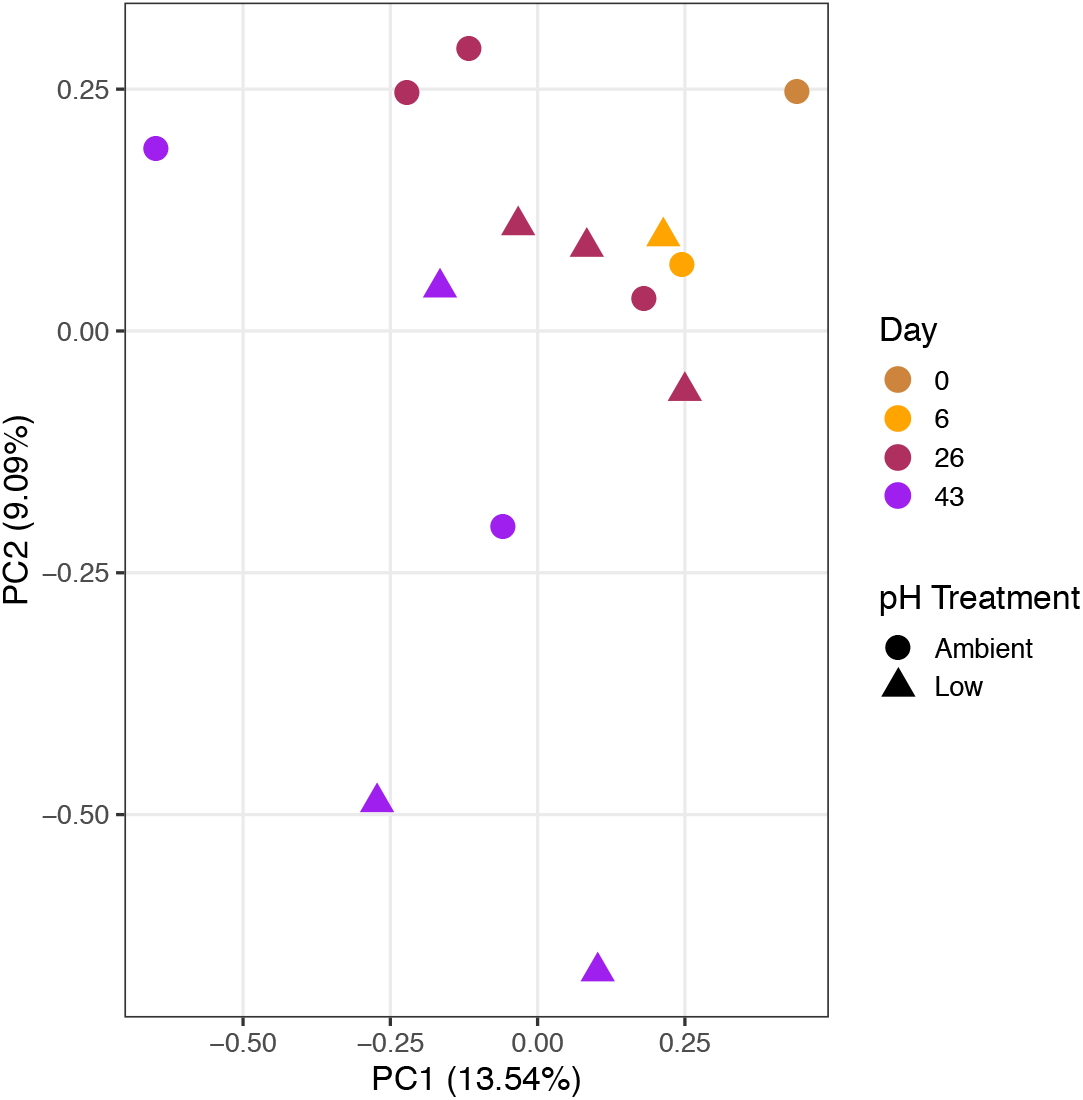
Principal component analysis of allele frequency data from larval samples collected throughout the course of the experiment. Allele frequency from 29,400 SNP’s were used for PCA. Sample color corresponds to day and sample shape corresponds treatment condition.

We identified SNPs that changed significantly in frequency (*i.e.*, outlier SNPs) between the day 0 larval population and the larvae sampled on day 6, 26, and 43 in each treatment. Outlier SNPs were identified using a rank-based approach and the observed allele frequency shift probabilities generated from the Fisher’s Exact and Cochran-Mantel-Haenszel tests. This analysis indicated pervasive signatures of selection in both treatments, with thousands of SNPs significantly changing in frequency throughout the course of the experiment relative to day 0 (Fig 4a). As the larvae sampled on day 6 were drawn from different replicate buckets as those sampled on days 26 and 43 (Fig. 1), outlier SNPs observed on all three sampling days point to candidate loci that may be putatively under selection in each pH environment. We used these robust SNPs to generate lists of pH-specific loci or overlapping loci (genes containing outlier SNPs that were responsive in each treatment). In total, we identified 99 ambient pH-specific loci 31 annotated), 88 low pH specific-loci (29 annotated), and 63 shared loci (24 annotated) based on transcriptome provided in Moreira *et al.* (2015)^30^ (see Supplementary File 1). Therefore, 58% of the loci exhibiting signatures of selection in the low pH treatment were unresponsive, and putatively neutral, in the ambient treatment (Fig. 6a). This finding provided an initial indication of the presence of cryptic variation in the population that may facilitate adaptation to low pH conditions.

**Figure 4.**
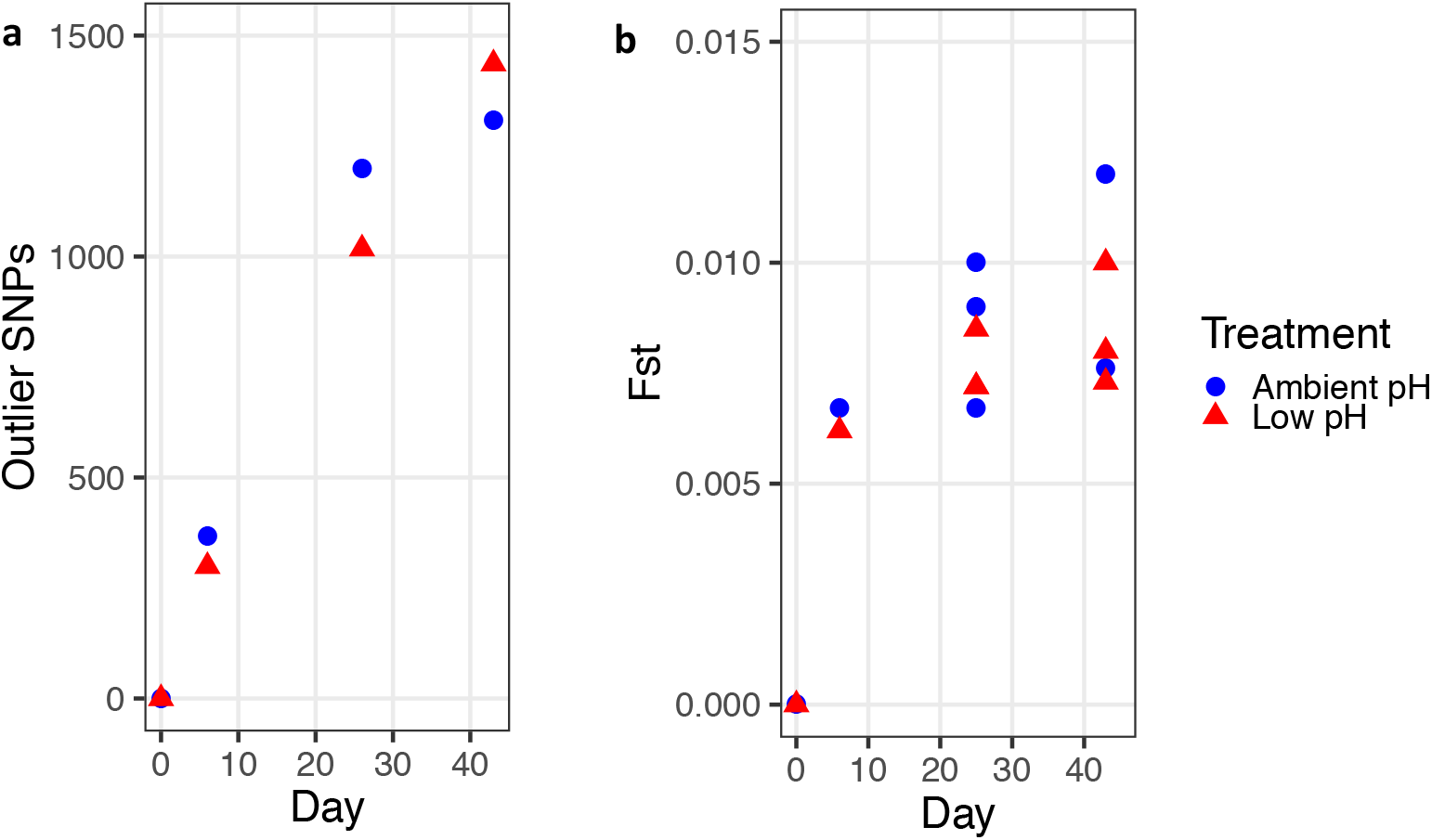
Changes in outlier SNPs and F_ST_ throughout early development. **(a)** The number of outlier SNPs identified in ambient and low pH treatment throughout the experiment. The number of outliers reported was standardized by the number of replicate buckets sampled on each day in order to account for increased power associated with increased replication **(b)** F_ST_ between the Day 0 larval population and the larval population in each treatment through day 43.

Another statistical metric of genetic differentiation, F_ST_, was used to identify changes in the magnitude of selection throughout development. We computed exome-wide (global) estimates of F_ST_ pairwise between the day 0 larval population and each available replicate bucket on all sampling days. The greatest change in F_ST_ occurred between day 0 and 6, before elevating more slowly thereafter, suggesting that the majority of selective mortality in *M. galloprovincialis* larvae occurred prior to day 6 (Fig. 4b).

### Size Separation

The size-separation of larvae on day 6 isolated the largest 18% from the smallest 82% in the ambient and the largest 21% from the smallest 79% in the low pH treatment (Fig. S1). Hereafter, these groups will be referred to as the fastest and slowest growers, respectively. Shell size on day 6 was significantly affected by treatment and size class (Linear Mixed Effects Model, p < 0.001). PCA using allele frequency data from the day 0 starting larval population and larval samples collected on day 6 revealed a strong genetic signature of size class (Fig. 5). Specifically, the fastest growers segregated along PC1 from the slowest growers in both treatments, with the day 0 larval population and day 6 larval population samples (from each treatment) falling in between the size-separated groups. The number of outlier SNP’s differentiating the fastest and slowest growers in each treatment, hereafter referred to as size-selected SNPs, was comparable: 963 outlier SNPs identified ambient and 846 outlier SNPs identified in the low pH treatment (outliers identified using Cochran-Mantel-Haenszel test). This led to the identification of 611 size-selected loci that were unique to the ambient pH treatment (225 annotated), 499 size-selected loci that were unique to the low pH treatment (184 annotated), and 154 size-selected loci (51 annotated) that were shared between environments^30^ (Supplementary File 2). Therefore, 76% of loci associated with fast shell growth in low pH were not associated with fast growth in the ambient treatment, and indicated that the fastest growers in the low pH environment come from a largely distinct genetic background (Fig 6b). F_ST_ analysis corroborated this trend, as elevated signatures of differentiation between the fastest growers in ambient and low pH, relative to differentiation between the slowest growers in ambient and low pH treatments as well as the entire day 6 larval populations in ambient and low treatments, were observed (Fig. S2). Thus, the size separation, in concert with the pH-specific signatures of selection observed throughout the entire larval period, further indicated a unique genetic background associated with fitness in the low pH treatment (Fig. 6).

**Figure 5.**
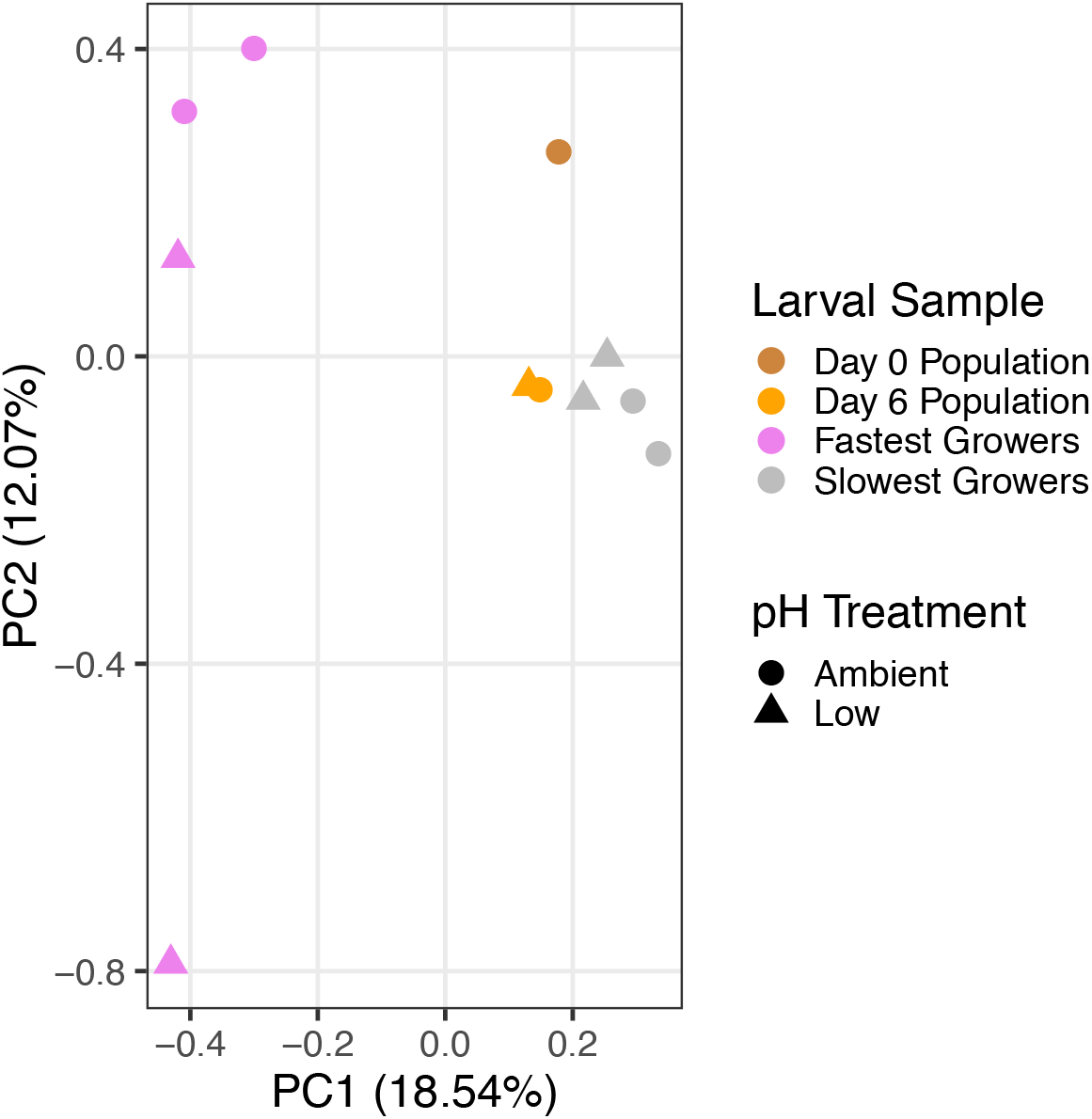
Principle component analysis of SNPs examining the genomic signature of shell growth. Allele frequency data using 29,400 SNPs identified in larval smples collected on day 6, 6, and size separated larvae from day 6 (i.e. fastest and slowest growers).

**Figure 6.**
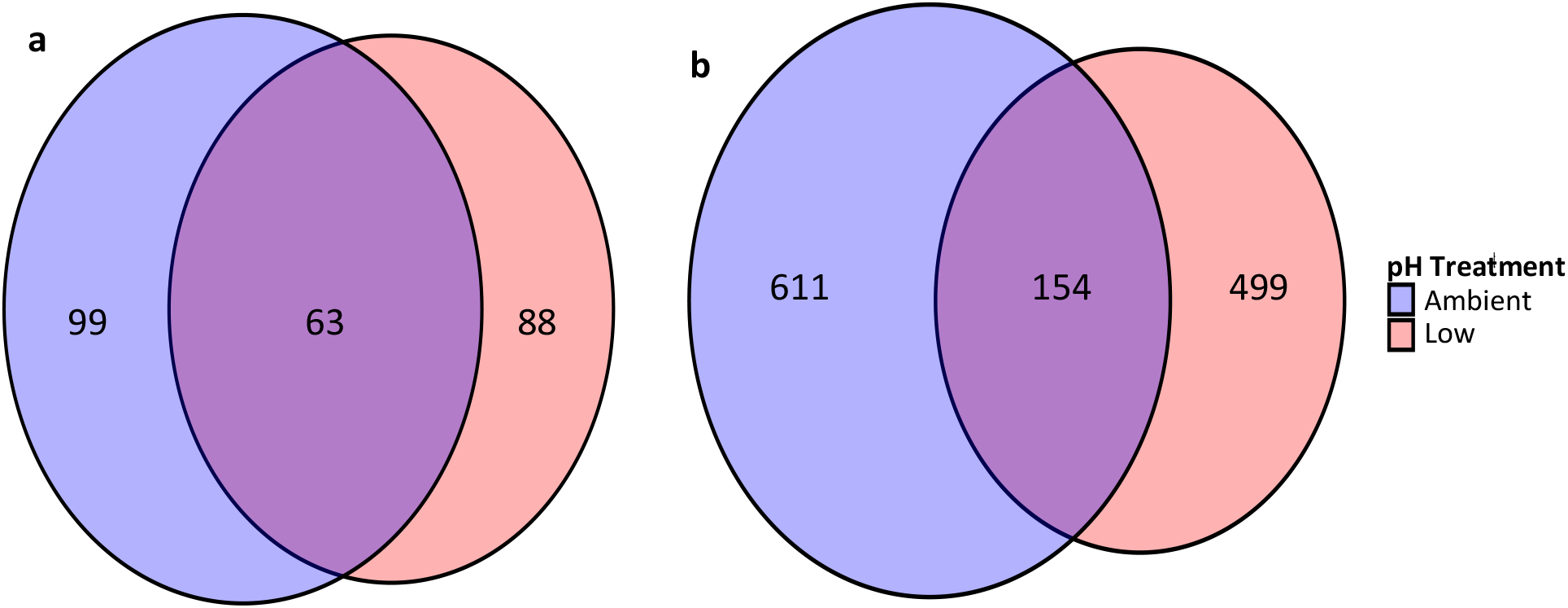
Genotype-environment interactions during selection in alternate pH environments. Venn diagrams show the extent of overlap between candidate outlier loci in ambient and low pH conditions **(a)** throughout the entire larval period (58% unique outlier loci in low pH treatment), and **(b)** between the fastest and slowest growers on day 6 (76% unique outlier loci in low pH treatment)

## Discussion

### Concurrent shifts in larval size and genetic variation throughout development

While previous work has shown strong negative effects of low pH on larval development in bivalves^25,27^, the results presented here suggest that standing variation within the species could facilitate rapid adaptation to ocean acidification. Observed shell length differences on days 3 and 7 matched expectations for the species based on previous work (−1 μm per 0.1 unit decrease in pH)^29^. However, this difference was reduced ∼50% by day 14. Mechanistically, low pH treatment effects on bivalve larval shell growth are driven by the limited capacity of larvae to regulate carbonate chemistry, specifically aragonite saturation state, in their calcifying space^31–33^. Our data show that this physiological limitation is greatest prior to day 7, after which low pH larvae were able to partially recover in size compared to larvae reared in the ambient treatment.

It is likely that partial recovery of shell size in low pH larvae observed by day 14 was, at least in part, driven by natural selection. We have previously shown that the smallest D-veligers in the low pH treatment display an increased prevalence of morphological abnormalities, which likely become lethal during the shell growth period^29^. Directional selection against this phenotypic group would shift the size distribution closer to that of larvae reared in ambient conditions, as we observed. The unique genetic backgrounds of the fastest growing larvae in low pH at 6 days post-fertilization, as well as the unique outlier loci identified in the low pH environment throughout the larval period, further strengthen the notion that these phenotypic trends were rooted in changes in the larval population’s underlying genetic variation.

Increasing genetic differentiation through time, as evidenced by PCA, outlier loci identification, and F_ST_ in both treatments further suggested the process of selection during the shell growth period. However, the observed trends in F_ST_ highlighted a developmental point of heightened selection, even before larval size distributions began to converge. Specifically, when F_ST_ is scaled by duration of treatment exposure, the genomic differentiation between the day 0 larval population and the larval population on day 6 was three and five times greater than that observed between the day 0 larval population and the larval population on days 26 and day 43, respectively. This suggests that a largely singular, intense selection event occurred prior to day 6 and may be responsible for the majority of genetic differentiation that occurs during larval growth and settlement. We recently identified two specific early developmental processes that are sensitive to low pH conditions and occur in this timeframe^29^. These processes include the formation of the shell field (early trochophore stage) and the transition between growth of first and second larval shell (late trochophore stage), both of which occur within 48 hours of fertilization, resulting in a suite of size-dependent morphological abnormalities that likely become lethal during the shell growth period^29^. Traditionally, metamorphosis from the swimming D-veliger to the settled juvenile is regarded as the main genetic bottleneck during the development of marine bivalve larvae^34^. Our sampling from the embryo stage through settlement, however, suggests that there is a major selection event prior to day 6 that may have an even larger effect on shaping genotypes of settled juveniles any selection thereafter.

Additional factors that may have allowed the larvae reared in led to the observed phenotypic dynamics are food-augmented acclimation and selective mortality via food competition. It has been demonstrated that increased energy availability can allow marine invertebrates to withstand pH stress^35^ and, in the case of *Mytilus edulis*, food availability can mitigate the negative effects of ocean acidification^36^. This compensation, however, is unlikely in our experiment. Our algal concentrations during the period of phenotypic convergence (days 7-26) fell below optimal concentrations reported for the species^37,38^. The substantial delay in the duration of the pelagic phase relative to published developmental timelines for the species^38,39^, further demonstrates that the larvae were indeed food limited in each treatment. This food limitation may have induced intraspecific competition and facilitated the selective mortality of less fit genotypes, thereby producing the pervasive signatures of selection observed in both treatments starting on day 6. Selection via differential mortality may have been concentrated on the smallest larvae in low pH, thus driving the phenotypic convergence between treatments. Ultimately, surviving larvae in the low pH treatment were able to partially compensate for the negative effect of CO_2_-acidification on calcification kinetics^32^, though the extent to which multi-generational selection would allow the population to completely recover the offset in shell size is an important area of future research. These results bolster emerging studies that have demonstrated the importance of standing variation in shaping the pH tolerance pf coastal marine species^40–42^.

### The role of cryptic genetic variation in extreme low pH adaptation

We identified hundreds of loci responding to each pH treatment throughout the larval period. Notably, 58% of our candidate low pH loci were statistically unchanged in the ambient conditions. While some of this treatment disparity may be an artifact (*i.e.* false positives in the low pH or false negatives in the ambient treatment), it is unlikely that this is the case for all the unique SNPs identified. These data thus provide evidence that loci that were not critical for fitness in the ambient environment came under strong selection in the low pH treatment. Associating shell growth to its underlying genetic background in each environment strengthened this conclusion. Specifically, there was an exceedingly small amount of overlap in size-selected loci between treatments (76% of size-selected loci were unique to the low pH environment) and a relative elevation of F_ST_ differentiation between the fastest growers from the low and ambient pH buckets. These patterns display a classic genotype-environment interaction in which a particular genetic background exhibits a specific trait value in one environment (e.g. accelerated shell growth in ambient pH), while an alternate genetic background leads to the same trait value when the environment shifts^14^. As shell growth is a direct proxy for fitness^29,43^, these data suggest that the most fit genotypes in ambient conditions may not be the individuals that harbor the adaptive genetic variation necessary to improvs fitness in simulated ocean acidification.

Ultimately, our observation of genotype-environment interactions in response to a dramatic shift in seawater pH provides strong evidence that adaptation to ocean acidification may be fueled by cryptic genetic variation. The role of cryptic genetic variation during adaptation to novel and extreme environmental conditions is becoming increasingly recognized^12,14,15,44,45^, though its relative importance has yet to be demonstrated as clearly in nature as it has been in theory^9,14,16^. Our data not only suggest the role of cryptic genetic variation in rapid adaptation, but also demonstrate this phenomenon in the context of a non-model species subject to global change. The economic and ecological importance of marine mussels, as well as their global exposure to declining seawater pH, highlights the need to conserve standing variation in order to allow the adaptive capacity of natural populations to play out as climate change progresses.

Many outlier loci in low pH were also outliers in the ambient treatment (42%). This likely represents the action of selection against recessive homozygotes within the population, termed genetic load^46^, and selection by the laboratory regime. The influence of genetic load has been demonstrated to induce signatures of selection in “neutral” environments in a range of highly fecund species, such as plants and marine bivalves^34^. This was likely amplified in the present study by our crossing scheme, which purposefully induced equal proportions of all pairwise crosses, thereby maximizing the likelihood of lethal, or less “fit”, homozygotes in the day 0 larval population. These shared signatures of selection could be further associated with selective pressures induced by the laboratory conditions, such as salinity, temperature, or the food resources, which are independent of the pH manipulation. While the laboratory conditions were designed to optimize larval growth and mirror environmental conditions at the collection site, it is unsurprising that a specific set of environmental variables (*i.e.* lab conditions) favors a subset of genetic backgrounds in a species exhibiting such high levels of heterozygosity^47^.

### Putative targets of selection as ocean acidification progresses

The low-pH specific loci we identified (loci with outlier SNPs in every replicate bucket and across all sampling days) provide targets of natural selection as ocean acidification progresses. These included a HSPA1A gene (Swissprot ID: Q8K0U4), which encodes heat shock protein 70 (HSP70), one of a group of gene products whose expression is induced by physiological stressors and generally work to mediate/prevent protein denaturation and folding^48^. While substantial evidence has documented the role of HSP70 in the thermal stress response across a range of taxa^48^, emerging transcriptomic studies have also demonstrated the protein’s role in the physiological response to low pH conditions in echinoderms^49^ and bivalves^50^. Another notable candidate locus is a tyrosinase-like protein (Swissprot ID: H2A0L0). Tyrosinase genes are known to influence biomineralization in marine bivalves during larval^51,52^ and juvenile stages^53^, a process that is fundamentally affected by changes in seawater chemistry. While these gene expression-based studies provided initial insight into the underlying physiological responses to changes in seawater chemistry in marine bivalves, our study demonstrates the presence of underlying genetic variation within these putative loci. This provides, to our knowledge, the first documentation of standing genetic variation at functionally relevant loci within marine bivalves, and ultimately offers robust evidence for the species’ capacity to adapt to extreme changes in seawater pH. We are currently investigating these candidates more intensely through a combination of comparative transcriptomics, quantitative PCR, and *in situ* hybridization in *M. galloprovincialis* larvae (Kapsenberg *et al*., unpublished).

### Conclusions

Species persistence as global climate change progresses will, in part, hinge upon their ability to evolve in response to the shifting abiotic environment^21^. Our data suggest that the economically and ecologically valuable marine mussel, *M. galloprovincialis*, currently has the standing variation necessary to adapt to ocean acidification, though with a potential trade-off of shell size. We have demonstrated that much of this variation may be cryptic in ambient pH conditions, yet bolsters a fitness-related trait (shell growth) when seawater pH is reduced. Ultimately, these findings support conservation efforts aimed at maintaining variation within natural populations to increase species resilience to future ocean conditions. In a broader evolutionary framework, the substantial levels of genetic variation present in natural populations have puzzled evolutionary biologists for decades^54^. Though this study does not address the processes that maintain this variation during periods of environmental stasis, we demonstrate its utility in rapid adaptation, thereby advancing our understanding of the mechanisms by which natural populations evolve to abrupt changes in the environment.

## Methods

### Larval cultures

Mature *M. galloprovincialis* individuals were collected in September 2017 from the underside of a floating dock in Thau Lagoon (43.415°N, 3.688°E), located in Séte, France. Thau Lagoon has a mean depth of 4 m and connection to the Mediterranean Sea by three narrow channels. pH variability at the collection site during spawning season ranges from pH_T_ 7.80 to 8.10^29^. Mussels were transported to the Laboratoire d’Océanographie (LOV) in Villefranche-sur-Mer, France and stored in a flow-through seawater system maintained at 15.2°C until spawning was induced.

Within 3 weeks of the adult mussel collection, individuals were cleaned of all epibiota using a metal brush, byssal threads were cut, and mussels were warmed in seawater heated to 27°C (∼+12°C of holding conditions) to induce spawning. Individuals that began showing signs of spawning were immediately isolated, and allowed to spawn in discrete vessels, which were periodically rinsed to remove any potential gamete contamination. Gametes were examined for viability and stored on ice (sperm) or at 16°C (eggs). In total, gametes from 12 females and 16 males were isolated to generate a genetically diverse starting larval population. To produce pairwise crosses, 150,000 eggs from each female were placed into sixteen separate vessels, corresponding to the sixteen founding males. Sperm from each male was then used to fertilize the eggs in the corresponding vessel, thus eliminating the potential effects of sperm competition and ensuring that every male fertilized each female’s eggs. After at least 90% of the eggs had progressed to a 4-cell stage, equal volumes from each vessel were pooled to generate the day 0 larval population (∼2 million individuals), from which the replicate culture buckets were seeded. 100,000 individuals were added to each culture buckets (N = 12, 18 embryos mL^−1^). The remaining embryos were frozen in liquid nitrogen, and stored at −80°C for DNA analysis of the day 0 larval population. Likewise, gill tissue was collected from all founding individuals and similarly stored for downstream DNA analyses. Larvae were reared at 17.2°C for 43 days. Starting on day 4, larvae were fed 1.6 × 10^8^ cells of *Tisochrysis lutea* daily. Beginning on day 23, to account for growth and supplement diet, larvae diet was complemented with 0.2 μL of 1800 Shellfish Diet (Reed Mariculture) (days 23-28 and day 38) and approximately 1.6 × 10^8^ cells of *Chaetoceros gracilis* (days 29-37 and 39-41).

### Larval sampling

We strategically sampled larvae throughout the experiment to observe phenotypic and genetic dynamics across key developmental events, including the trochophore to D-veliger transition (day 6), the shell growth period (day 26), and the metamorphosis from D-veligers to settlement (day 43). On day 6 of the experiment, larvae were sampled from three of the six replicate buckets per treatment. A subset of larvae (N = 91-172) from each bucket was isolated to obtain shell length distributions of larvae reared in the two treatments. The remaining larvae (∼25,000) were separated by shell size using a series of six Nitex mesh filters (70 μm, 65 μm, 60 μm, 55 μm, 50 μm, and 20 μm; Figure S1) and frozen at −80°C. The smallest size group contained larvae arrested at the trochophore stage, and therefore unlikely to survive. The remaining five size classes isolated D-veligers from the smallest to the largest size. The shell length distribution of the larvae was used to inform, *a posteriori*, which combination of size classes would produce groups of the top 20% and bottom 80% of shell sizes from each treatment. The relevant size groups from two replicates per treatment were then pooled for downstream DNA analysis of each phenotypic group. For the third replicate, *a posteriori*, all size groups were pooled in order to compute the allele frequency distribution from the entire larval population in each treatment on day 6. This sample was incorporated into analyses of remaining replicate buckets, which were specifically used to track shifts in phenotypic and genetic dynamics throughout the remainder of the larval period in each treatment.

Following size separation on day 6, the remaining replicate buckets (N = 3 per treatment) were utilized to track changing phenotypic and allele frequency distributions in the larval population through settlement. Larvae were sampled for size measurements on day 3 (n = 30-36), day 7 (N = 38-71), day 14 (N = 37-104), and day 26 (N = 49-112). Also on day 26, an additional ∼1,000 larvae per replicate were frozen and stored at −80°C pending DNA analysis. Finally, on day 43, settled individuals were sampled from each bucket (settlement was first observed on day 40 in all buckets). Treatment water was removed, and culture buckets were washed three times with FSW to remove unsettled larvae. Individuals that remained attached to the walls of the bucket were frozen and stored at −80°C for DNA analysis.

### Culture system and seawater chemistry

Larvae were reared in a temperature-controlled sea table (17.2°C) and 0.35 μm filtered and UV-sterilized seawater (FSW), pumped from 5 m depth in the bay of Villefranche. Two culture systems were used consecutively to rear the larvae, both of which utilized the additions of pure CO_2_ gas for acidification of FSW. First, from days 0-26 the larvae were kept in a flow-through seawater pH-manipulation system described in Kapsenberg *et al.* (2017)^55^. Briefly, seawater pH (pH_T_ 8.05 and pH_T_ 7.4) was controlled in four header tanks using a glass pH electrode feedback system (IKS aquastar) and pure CO_2_ gas addition and constant CO_2_-free air aeration. Two header tanks were used per treatment to account for potential header tank effects. Each header tank supplied water to three replicate culture buckets (drip rate of 2 L h^−1^), fitted with a motorized paddle and Honeywell Durafet pH sensors for treatment monitoring (see Kapsenberg et al 2017 for calibration methods).

On day 27 of the experiment, the flow-through system was stopped due to logistical constraints and treatment conditions were maintained, in the same culture buckets, using water changes every other day. For water changes 5 L of treatment seawater (70% of total volume) was replaced in each culture using FSW pre-adjusted to the desired pH treatment. Seawater pH in each culture bucket was measured daily, and before and after each water change.

All pH measurements (calibration of Durafets used from day 0-26 and monitoring of static cultures from day 27-43) were conducted using the spectrophotometric method and purified *m*-cresol dye and reported on the total scale (pH_T_)^56^. Samples for total alkalinity (*A*_T_) and salinity were taken from the header tanks every 2-3 days from days 0-26 and daily during the remainder of the experiment. *A*_T_ was measured using an open cell titration on Metrohhm Titrando 888^56^. Accuracy of *A*_*T*_ measurements was determined using comparison to a certified reference material (Batch #151, A. Dickson, Scripps Institution of Oceanography) and ranged between −0.87 and 5.3 μmol kg^−1^, while precision was 1.23 μmol kg^−1^ (based on replicated samples, *n* = 21). Aragonite saturation and *p*CO2 were calculated using pH and *A*_*T*_ measurements and the *seacarb* package in R^57^ with dissociation constants K_1_ and K_2_ ^58^, Kf^59^ and Ks^60^. Seawater chemistry results are presented in the electronic supplementary, Tables S1–S2.

### Shell Size Analysis

Shell size was determined as the maximum shell length parallel to the hinge using brightfield microscopy and image analysis in ImageJ software. All statistical analyses were conducted in R (v. 3.5.3)^61^. As larval shell length data did not pass normality tests (Shapiro-Wilk test), shell-size was log-transformed to allow parametric statistical analysis. We tested the effect of day, treatment, and the interaction of the two using linear mixed effects models, with day and treatment as fixed effects and replicate bucket as a random effect (*lmer*). Effects of treatment and size class on log-transformed shell length from size-separated larvae were also analyzed using a linear-mixed effect model. Size class, treatment, and their interaction were fixed effects, while larval bucket was a random effect.

### DNA extraction and exome sequencing

We implemented exome capture, a reduced-representation sequencing approach, to identify SNPs and their frequency dynamics throughout the course of the experiment. Exome capture targets the protein-coding region of the genome, and thus increases the likelihood that identified polymorphisms have functional consequences^62^. Genomic DNA from each founding individual and larval sample was extracted using the EZNA Mollusc Extraction Kit, according to manufacturer’s protocol. DNA was quantified with a Qubit, and quality was determined using agarose gel, Nanodrop (260/280), and TapeStation analysis.

Genomic DNA was hybridized to a customized exome capture array designed and manufactured by Arbor Biosciences (Ann Arbor, Michigan) and using the species transcriptome provided in Moreira et al. (2015). Specifically, in order to design a bait set appropriate for capture of genomic DNA fragments, 90-nucleotide probe candidates were tilled every 20 nucleotides across the target transcriptome contigs. These densely-tiled candidates were MEGABLASTed to the *Mytilus galloprovincialis* draft genome contigs available at NCBI (GCAA_001676915.1_ASM167691v1_genomic.fna), which winnowed the candidate list to only baits with detected hits of 80 nucleotides or longer. After predicting the hybrid melting temperatures for each near-full-length hit, baits were further winnowed to those with at most two hybrids of 60°C or greater estimated melting temperature in the *M. galloprovincialis* genome. This collection of highly specific baits with near-full-length hits to the draft genome were then down-sampled to a density of roughly one bait per 1.9 kbp of the final potential target space, in order to broadly sample the target while still fitting within our desired number of myBaits kit oligo limit. The final bait set comprises 100,087 oligo sequences, targeting 94,668 of the original 121,572 transcript contigs.

Genomic DNA from each sample was subject to standard mass estimation quality control, followed by sonication using a QSonica QR800 instrument and SPRI-based dual size-selection to a target modal fragment length of 350 nucleotides. Following quantification, 300 ng total genomic DNA was taken to library preparation using standard Illumina Truseq-style end repair and adapter ligation chemistry, followed by six cycles of indexing amplification using unique eight nucleotide dual index primer pairs. For target enrichment with the custom myBaits kit, 100 ng of each founder-derived library were combined into two pools of 14 libraries each, whereas 450 ng of each embryonic and larval-pool derived library were used in individual reactions. After drying the pools or individual samples using vacuum centrifugation to 7 μL each, Arbor followed the myBaits procedure (v. 4) using the default conditions and overnight incubation to enrich the libraries using the custom probe set. After reaction cleanup, half (15 μL) of each bead-bound enriched library was taken to standard library amplification for 10 cycles using Kapa HiFi polymerase. Following reaction cleanup with SPRI, each enriched library or library pool was quantified using qPCR, indicating yields between 30 and 254 ng each.

The captured libraries were sequenced at the University of Chicago Genomics Core Facility on three lanes of Illumina HiSeq 4000 using 150-bp, paired-end reads. The captured adult libraries were sequenced on an individual lane, while the twenty-two, pooled larval samples were split randomly between the remaining two lanes. Average coverage for founding individuals was 40x, while average coverage in pooled samples was 100x.

### Read Trimming and Variant Calling

Raw DNA reads were filtered and trimmed using Trimmomatic^63^ and aligned to the species reference transcriptome provided in Moreira *et al.* (2015) using bowtie2^64^. Variants in the founding individuals were identified using the Genome Analysis Toolkit’s^65^ Unified Genotyper. These variants were filtered using VCFTOOLs^66^ with the following specifications: Minor Allele Frequency of 0.05, Minimum Depth of 10x, and a Maximum Variant Missing of 0.75. The resulting .vcf files provided a list of candidate bi-allelic polymorphisms to track at each time point, treatment, and phenotypic group in the larval samples. Accordingly, GATK’s Haplotype Caller was used to identify these candidate polymorphisms within each larval alignment file, and the resulting .vcf was filtered using VCFTOOLs and the following specifications: Minor Allele Frequency of 0.01, Minimum Depth of 50x, and Maximum Depth of 450x. Only variants that passed quality filtering and were identified in all larval samples (i.e. each day, treatment, and phenotypic group) were retained for downstream analyses. This process resulted in a candidate SNP list of 29,400 variants. Allele frequencies for each variant were computed as the alternate allelic depth divided by total coverage at the locus.

### Allele Frequency Analysis

To explore how the allele frequency of the 29,400 SNPs changed in each environment throughout the course of the experiment, we used a combination of PCA, outlier loci identification tests, and a statistical test of genomic differentiation (F_ST_). We visualized patterns of genetic variation throughout the experiment with PCA (*prcomp* function in R). Prior to PCA, the allele frequency matrix (the generation of which is provided in the previous section) was centered and scaled using the *scale* function in R. Only larval samples that encompassed the full phenotypic distribution within a particular bucket were included in this analysis. In other words, the rows of the allele frequency matrix corresponding to larval samples that were selectively segregated based on shell size were removed, and PCA was run using the day 0 larval population and larval samples collected from each treatment on days 6, 26, and 43. A separate PCA was then implemented using data from the day 0 larval population and all day 6 larval samples, which included discrete size groups from each treatment. This analysis thus explicitly examined a genomic signature of the individuals that were phenotypically distinct.

We next sought to identify the presence, number, and treatment-level overlap of genetic variants that significantly changed in frequency between larval samples. Specifically, Fisher’s Exact test (FET) and the Cochran-Mantel-Haenszel (CMH) test were used to generate probabilities of observed allele frequency changes, using the package *Popoolation*^67^ in R. P-values for each SNP were converted to q-values in the R package *qvalue*^68^, and outliers were identified as those SNPs with a q-value <0.01. The FET was used to identify outliers between the day 0 larval population and the day 6 larval populations in each treatment (no treatment replicates were available for this comparison). For all remaining allele frequency comparisons (in which replicate bucket information was available), the CMH test was used to identify consistent allele frequency shifts among replicate buckets. We used this test to identify significant allele frequency changes between the day 0 larval population and the day 26 treatment replicates (N = 3), the day 0 larval population and the settled individuals treatment replicates (ambient pH treatment: N = 2, low pH treatment: N = 3), and between the top 20% and bottom 80% of growers in each treatment (N = 2).

To provide a third, independent metric of genomic change in the larval population throughout the experiment, we computed the F_ST_ statistic for a series of comparisons. Specifically, we implemented a methods-of-moments estimator of F_ST_ from Pool-seq data in an analysis of variance framework, as described in Hivert et al. (2018)^69^ (*poolFstat* package). A global (exome-wide) F_ST_ statistic was computed pairwise between the day 0 larval population and the day 6 ambient and low pH larvae replicate buckets, day 26 ambient and low pH larvae replicate buckets, and settled individuals from all replicate buckets in ambient and low pH. F_ST_ was also computed to compare differentiation between phenotypic groups (top 20% and bottom 80% of growers) on day 6. Estimates of pool sizes were based on the known day 0 larval population size, the known volume of larvae sampled, and the observed declines in larval culture density. Input pool sizes were as follows: 100,000 for day 0 larval population; 25,000 for ambient and low pH larval samples on day 6; 5,250 for top 20% of growers in ambient conditions on day 6; 4,250 for top 20% of growers in low pH conditions on day 6; 19,750 for bottom 80% of growers in ambient conditions 20,750 for bottom 80% of growers in low pH conditions on day 6; 2,500 larval samples on day 26; 250 for settled individuals collected on day 43. Equal densities within each treatment were assumed as no discernable difference in mortality between treatments was observed (MCB, *pers. obs.*).

### Gene Identification/Ontologies

We next sought to explore the biological pathways that were associated with survivorship in each pH treatment and/or size group during the experiment. To accomplish this, we indexed the outlier loci that contained variants with significant frequency changes using the annotated transcriptome provided in Moreira *et al.* (2015). Their annotation utilized NCBI’s nucleotide and non-redundant, Swissprot, KEGG, and COG databases, thus providing a thorough survey of potential genes and pathways associated with our candidate SNPs. We generated gene lists for pH-specific loci, which were identified as loci that showed signatures of selection on all sampling days and were unique to each environment. We also generated a candidate gene list for loci that exhibited shared signatures of selection in each treatment. These lists thus only contain robust candidate loci (loci identified as outliers in multiple independent replicates), with potentially strong effect sizes (loci identified as outliers at multiple developmental stages). Lastly, we used the Moreira *et al.* (2015) annotation to explore the genes that exhibited signatures of selection for shell growth in ambient and low pH conditions, as well as shared signatures of selection for shell size in each treatment.

## Acknowledgements

We thank Samir Alliouane for extensive technical assistance during the completion of the experiment. We would also like to thank Angelica Miglioli for experimental assistance, Régis Lasbleiz for microalgal supply for larval feeding, Jacob Enk at Arbor Biosciences for guidance during exome capture, and the University of Chicago Genomics Core Facility for sequencing assistance. Lastly, we thank D. Rice and T. Price for insightful comments on the manuscript.

## Author Contributions

MCB conceived and designed the experiment with inputs from LK, JPG, and CAP. MCB and LK performed the experiment. MCB completed molecular lab work, with exome capture and sequencing assistance from Arbor Biosciences and the University of Chicago Genomics Core. MCB completed all bioinformatics, statistical, and computational analyses. MCB wrote the manuscript with inputs from LK, JPG, and CAP.

## Supplemental Information

**Figure S1.**
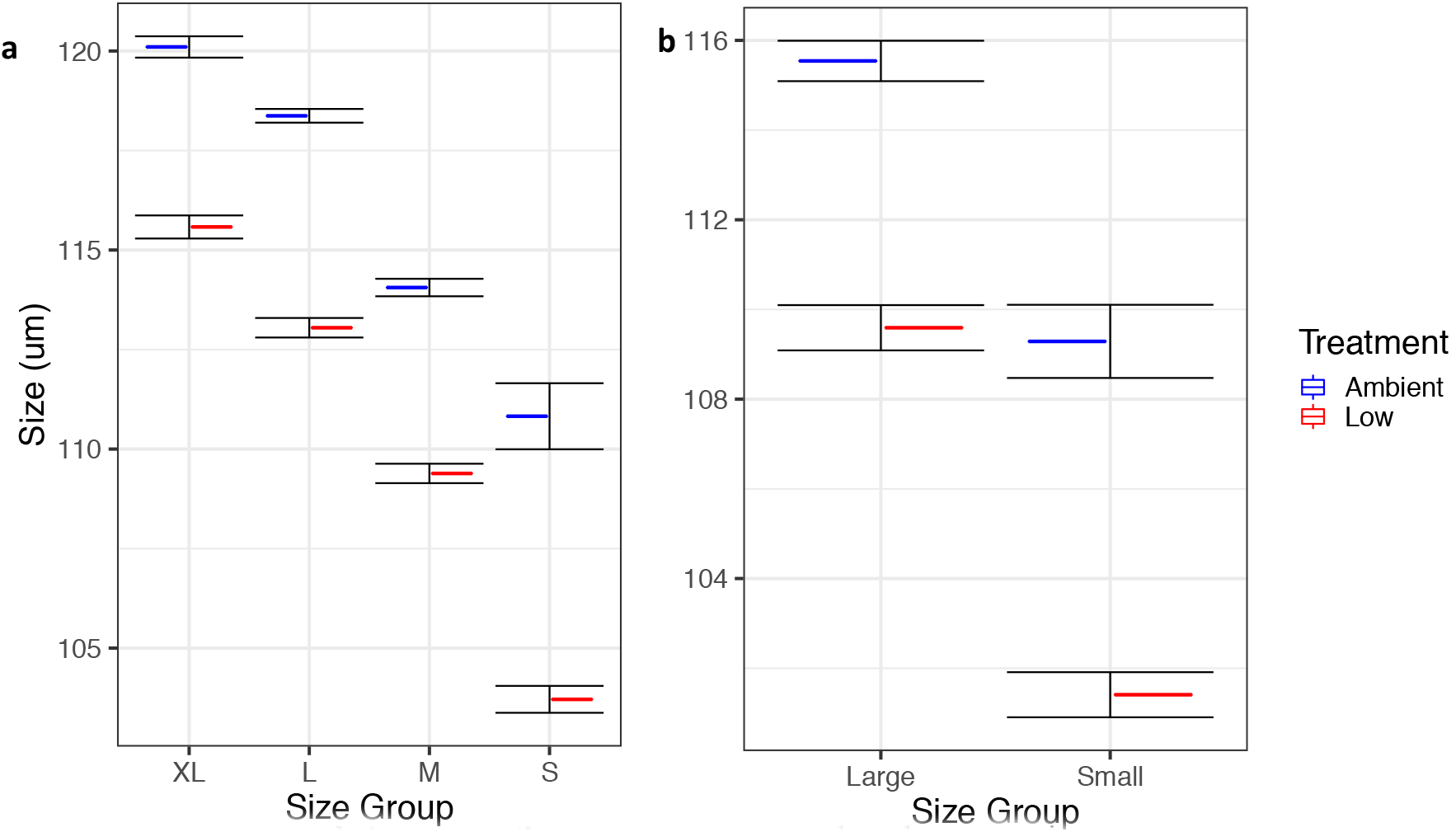
Efficacy of larval size separation method. **(a)** Pilot data demonstrating the selective isolation of 4 consecutively smaller groups of larvae (mean shell length +/− SE) in ambient and low pH cultured larvae (*N* = 53-324 per size group; XL = extra large; L = large; M = medium; S = small). **(b)** Shell length of larvae (mean +/− SE) in largest (top 20 %) and smallest (bottom 80 %) individuals in ambient and low pH conditions from present study (*N* = 62 – 177 per size group).

**Figure S2.**
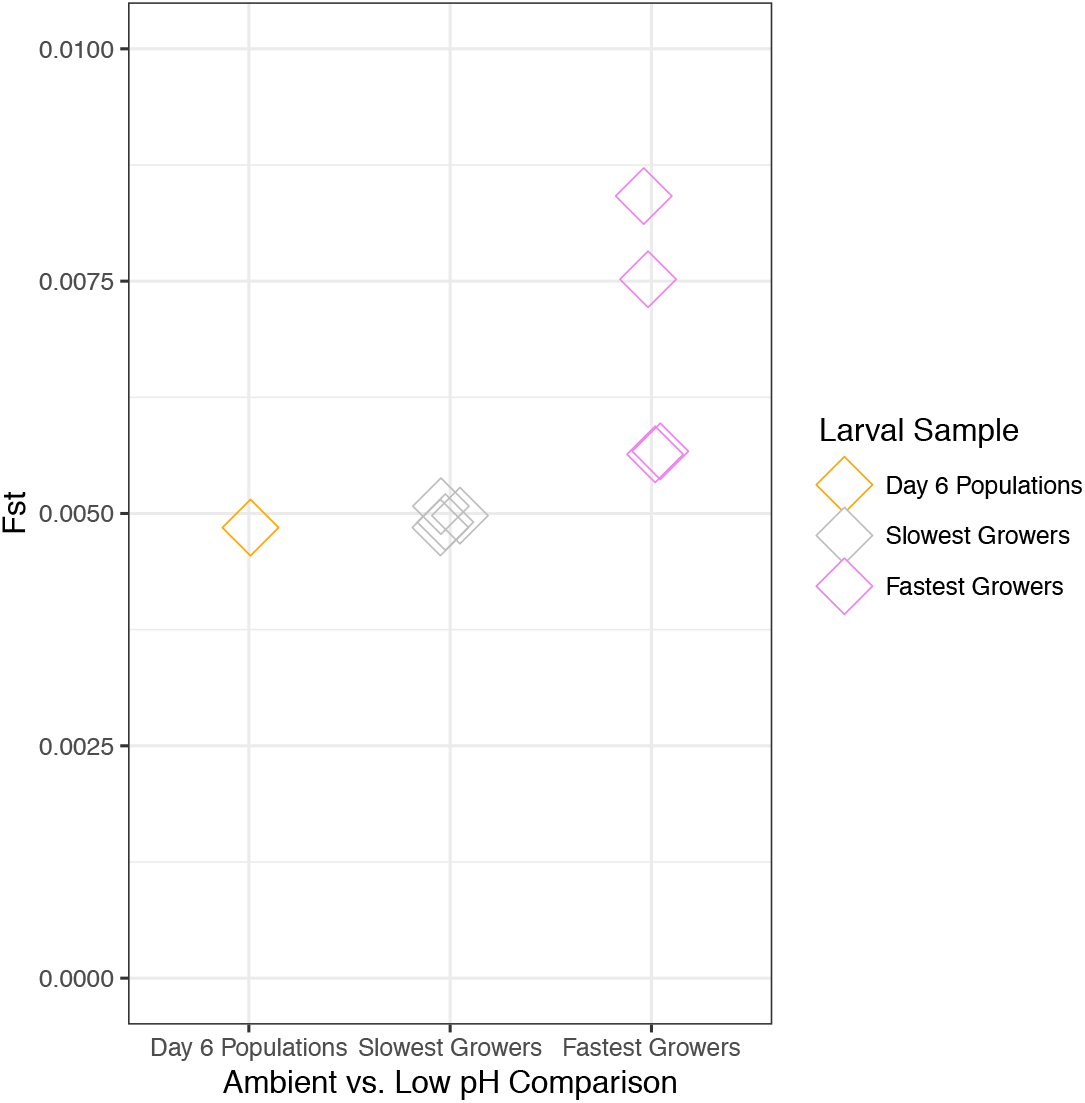
Genomic differentiation of fastest and slowest growers in ambient and low pH. Exome wide F_ST_ computed pairwise between ambient and low pH replicate buckets for the entire larval population and the size selected larvae isolated on day 6 (Day 6 Populations: *N*=1, Slowest and Fastest Growers comparisons: *N* = 4 pairwise comparisons).

**Table S1.** Carbonate chemistry for flow-through experimental system used on days 0-26, during which each of four header tanks (two per treatment) each distributed pH adjusted seawater to three replicate buckets. Low/Amb_1/2 correspond to treatment replicates drawing water from separate header tanks, thus one replicate bucket per header tank is represented in the table. Time-series pH and temperature data were generated using in autonomous sensor were generated in each representative replicate bucket, and averge values (+/− SD) are presented. Aragonite saturation (Ω_a_) and *p*CO_2_ were computed using average pH and AT for each representative replicate. Alkalinity (AT) and salinity samples were generated from discrete samples taken from each of the four header tanks every other day.

| pH treatment | pH<sub>T</sub> | $\Omega_a$ | $p\text{CO}_2$<br>($\mu\text{atm}$) | AT ($\mu\text{mol/kg}$) | T ($^{\circ}\text{C}$) | SAL (ppt) |
| --- | --- | --- | --- | --- | --- | --- |
| Low_1 | 7.43 (+/- 0.03) | 0.85 | 2127 | 2565<br>(n = 11) | 17.3 (+/-0.1) | 37.6 (+/-0.1)<br>(n = 11) |
| Low_2 | 7.43 (+/- 0.03) | 0.85 | 2130 | 2569<br>(n = 11) | 17.2 (+/-0.1) | 37.6 (+/-0.1)<br>(n = 11) |
| Amb_1 | 8.01 (+/- 0.01) | 3.32 | 378 | 2565<br>(n = 11) | 17.2 (+/-0.1) | 37.6 (+/-0.1)<br>(n = 11) |
| Amb_2 | 8.01 (+/- 0.01) | 3.31 | 378 | 2561<br>(n = 11) | 17.2 (+/-0.1) | 37.6 (+/-0.1)<br>(n = 11) |

**Table S2.**
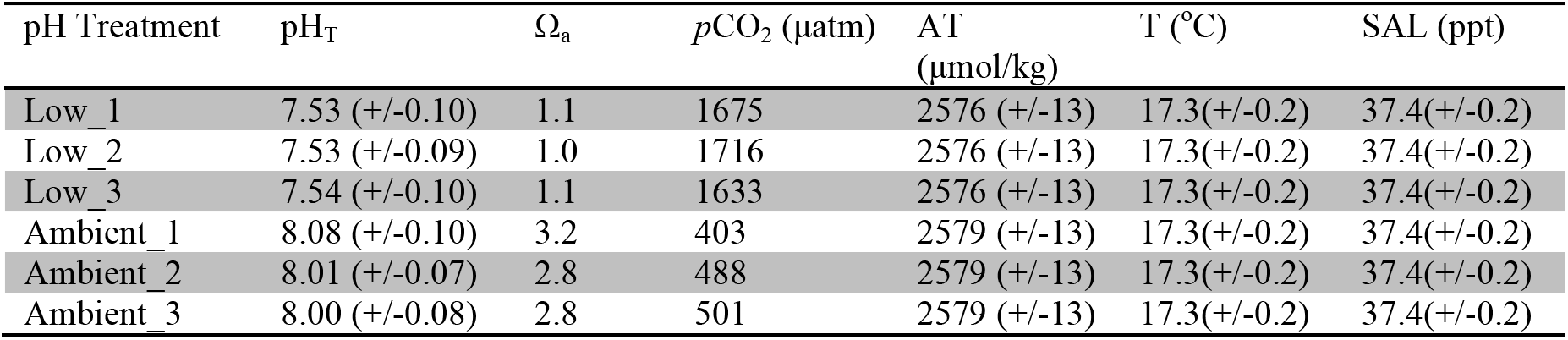
Carbonate chemistry generated from static cultures used to rear larvae from days 27-43. Low/Ambient_# correspond to each remaining replicate bucket during this portion of the experiment. Discrete measurements of pH, total alkalinity, temperature, and salinity were taken daily. Aragonite saturation (Ω_a_) and *p*CO_2_ were computed using average pH and AT for each replicate.

| pH Treatment | pH<sub>T</sub> | $\Omega_a$ | $p\text{CO}_2$ ($\mu\text{atm}$) | AT<br>($\mu\text{mol/kg}$) | T ($^{\circ}\text{C}$) | SAL (ppt) |
| --- | --- | --- | --- | --- | --- | --- |
| Low_1 | 7.53 (+/-0.10) | 1.1 | 1675 | 2576 (+/-13) | 17.3(+/-0.2) | 37.4(+/-0.2) |
| Low_2 | 7.53 (+/-0.09) | 1.0 | 1716 | 2576 (+/-13) | 17.3(+/-0.2) | 37.4(+/-0.2) |
| Low_3 | 7.54 (+/-0.10) | 1.1 | 1633 | 2576 (+/-13) | 17.3(+/-0.2) | 37.4(+/-0.2) |
| Ambient_1 | 8.08 (+/-0.10) | 3.2 | 403 | 2579 (+/-13) | 17.3(+/-0.2) | 37.4(+/-0.2) |
| Ambient_2 | 8.01 (+/-0.07) | 2.8 | 488 | 2579 (+/-13) | 17.3(+/-0.2) | 37.4(+/-0.2) |
| Ambient_3 | 8.00 (+/-0.08) | 2.8 | 501 | 2579 (+/-13) | 17.3(+/-0.2) | 37.4(+/-0.2) |

## References

1. Umina, P. A., Weeks, A. R., Kearney, M. R., McKechnie, S. W. & Hoffmann, A. A. A Rapid Shift in a Classic Clinal Pattern in Drosophila Reflecting Climate Change. Science 308, 691–693 (2005).

2. Rodríguez-Trelles, F., Tarrío, R. & Santos, M. Genome-wide evolutionary response to a heat wave in Drosophila. Biol. Lett. 9, 20130228 (2013).

3. Bergland, A. O., Behrman, E. L., O’Brien, K. R., Schmidt, P. S. & Petrov, D. A. Genomic Evidence of Rapid and Stable Adaptive Oscillations over Seasonal Time Scales in Drosophila. PLOS Genet. 10, e1004775 (2014).

4. Campbell-Staton, S. C. et al. Winter storms drive rapid phenotypic, regulatory, and genomic shifts in the green anole lizard. Science 357, 495–498 (2017).

5. Barrett, R. D. H. & Schluter, D. Adaptation from standing genetic variation. Trends Ecol. Evol. 23, 38–44 (2008).

6. Orr, H. A. The genetic theory of adaptation: a brief history. Nat. Rev. Genet. 6, 119–127 (2005).

7. Messer, P. W., Ellner, S. P. & Hairston, N. G. Can Population Genetics Adapt to Rapid Evolution? Trends Genet. 32, 408–418 (2016).

8. Via, S. & Lande, R. Genotype-Environment Interaction and the Evolution of Phenotypic Plasticity. Evolution 39, 505–522 (1985).

9. Hermisson, J. & Wagner, G. P. The Population Genetic Theory of Hidden Variation and Genetic Robustness. Genetics 168, 2271–2284 (2004).

10. Hartl, D. L. & Dykhuizen, D. E. Potential for selection among nearly neutral allozymes of 6-phosphogluconate dehydrogenase in Escherichia coli. Proc. Natl. Acad. Sci. 78, 6344–6348 (1981).

11. Vieira, C. et al. Genotype-Environment Interaction for Quantitative Trait Loci Affecting Life Span in Drosophila melanogaster. Genetics 154, 213–227 (2000).

12. McGuigan, K., Nishimura, N., Currey, M., Hurwit, D. & Cresko, W. A. Cryptic Genetic Variation and Body Size Evolution in Threespine Stickleback. Evolution 65, 1203–1211 (2011).

13. Hayden, E. J., Ferrada, E. & Wagner, A. Cryptic genetic variation promotes rapid evolutionary adaptation in an RNA enzyme. Nature 474, 92–95 (2011).

14. Paaby, A. B. & Rockman, M. V. Cryptic genetic variation: evolution’s hidden substrate. Nat. Rev. Genet. 15, 247–258 (2014).

15. Gibson, G. & Dworkin, I. Uncovering cryptic genetic variation. Nat. Rev. Genet. 5, 681–690 (2004).

16. Masel, J. Cryptic Genetic Variation Is Enriched for Potential Adaptations. Genetics 172, 1985–1991 (2006).

17. Dykhuizen, D. & Hartl, D. L. Selective neutrality of 6PGD allozymes in E. coli and the effects of genetic background. Genetics 96, 801–817 (1980).

18. Sangster, T. A. et al. HSP90-buffered genetic variation is common in Arabidopsis thaliana. Proc. Natl. Acad. Sci. 105, 2969–2974 (2008).

19. Queitsch, C., Sangster, T. A. & Lindquist, S. Hsp90 as a capacitor of phenotypic variation. Nature 417, 618–624 (2002).

20. Grant, P. R. et al. Evolution caused by extreme events. Phil Trans R Soc B 372, 20160146 (2017).

21. Hoffmann, A. A. & Sgrò, C. M. Climate change and evolutionary adaptation. Nature 470, 479–485 (2011).

22. Hoegh-Guldberg, O. et al. In: Climate Change 2014: Impacts, Adaptation, and Vulnerability. Part B: Regional Aspects. Contribution of Working Group II to the Fifth Assessment Report of the Intergovernmental Panel on Climate Change. (Cambridge University Press, Cambridge, United Kingdom and New York, NY, USA).

23. Hönisch, B. et al. The Geological Record of Ocean Acidification. Science 335, 1058–1063 (2012).

24. Kroeker, K. J. et al. Impacts of ocean acidification on marine organisms: quantifying sensitivities and interaction with warming. Glob. Change Biol. 19, 1884–1896 (2013).

25. Gazeau, F. et al. Impacts of ocean acidification on marine shelled molluscs. Mar. Biol. 160, 2207–2245 (2013).

26. Thomsen, J., Haynert, K., Wegner, K. M. & Melzner, F. Impact of seawater carbonate chemistry on the calcification of marine bivalves. Biogeosciences 12, 4209–4220 (2015).

27. Kurihara, H. Effects of CO2-driven ocean acidification on the early developmental stages of invertebrates. Mar. Ecol. Prog. Ser. 373, 275–284 (2008).

28. Ventura, A., Schulz, S. & Dupont, S. Maintained larval growth in mussel larvae exposed to acidified under-saturated seawater. Sci. Rep. 6, 23728 (2016).

29. Kapsenberg L. et al. Ocean pH fluctuations affect mussel larvae at key developmental transitions. Proc. R. Soc. B Biol. Sci. 285, 20182381 (2018).

30. Moreira, R. et al. RNA-Seq in Mytilus galloprovincialis: comparative transcriptomics and expression profiles among different tissues. BMC Genomics 16, 728 (2015).

31. Waldbusser, G. G. et al. A developmental and energetic basis linking larval oyster shell formation to acidification sensitivity. Geophys. Res. Lett. 40, 2171–2176 (2013).

32. Waldbusser, G. G. et al. Saturation-state sensitivity of marine bivalve larvae to ocean acidification. Nat. Clim. Change 5, 273–280 (2015).

33. Melzner, F., Thomsen, J., Ramesh, K., Hu, M. Y. & Bleich, M. Mussel larvae modify calcifying fluid carbonate chemistry to promote calcification. Nat. Commun. 8, 1709 (2017).

34. Launey, S. & Hedgecock, D. High Genetic Load in the Pacific Oyster Crassostrea gigas. Genetics 159, 255–265 (2001).

35. Pansch, C., Schaub, I., Havenhand, J. & Wahl, M. Habitat traits and food availability determine the response of marine invertebrates to ocean acidification. Glob. Change Biol. 20, 765–777 (2014).

36. Thomsen, J., Casties, I., Pansch, C., Körtzinger, A. & Melzner, F. Food availability outweighs ocean acidification effects in juvenile Mytilus edulis: laboratory and field experiments. Glob. Change Biol. 19, 1017–1027 (2013).

37. Phillips, N. E. Effects of Nutrition-Mediated Larval Condition on Juvenile Performance in a Marine Mussel. Ecology 83, 2562–2574 (2002).

38. Pettersen, A. K., Turchini, G. M., Jahangard, S., Ingram, B. A. & Sherman, C. D. H. Effects of different dietary microalgae on survival, growth, settlement and fatty acid composition of blue mussel (Mytilus galloprovincialis) larvae. Aquaculture 309, 115–124 (2010).

39. Carl, C., Poole, A. J., Williams, M. R. & Nys, R. de. Where to Settle—Settlement Preferences of Mytilus galloprovincialis and Choice of Habitat at a Micro Spatial Scale. PLOS ONE 7, e52358 (2012).

40. Kelly, M. W., Padilla-Gamiño, J. L. & Hofmann, G. E. Natural variation and the capacity to adapt to ocean acidification in the keystone sea urchin Strongylocentrotus purpuratus. Glob. Change Biol. 19, 2536–2546 (2013).

41. Pespeni, M. H. et al. Evolutionary change during experimental ocean acidification. Proc. Natl. Acad. Sci. 110, 6937–6942 (2013).

42. Thomsen, J. et al. Naturally acidified habitat selects for ocean acidification–tolerant mussels. Sci. Adv. 3, e1602411 (2017).

43. Allen, J. D. Size-Specific Predation on Marine Invertebrate Larvae. Biol. Bull. 214, 42–49 (2008).

44. Le Rouzic, A. & Carlborg, Ö. Evolutionary potential of hidden genetic variation. Trends Ecol. Evol. 23, 33–37 (2008).

45. McGuigan, K. & Sgrò, C. M. Evolutionary consequences of cryptic genetic variation. Trends Ecol. Evol. 24, 305–311 (2009).

46. Turner, J. R. G. & Williamson, M. H. Population Size, Natural Selection and the Genetic Load. Nature 218, 700 (1968).

47. Romiguier, J. et al. Comparative population genomics in animals uncovers the determinants of genetic diversity. Nature 515, 261–263 (2014).

48. Feder, M. E. & Hofmann, G. E. HEAT-SHOCK PROTEINS, MOLECULAR CHAPERONES, AND THE STRESS RESPONSE: Evolutionary and Ecological Physiology. Annu. Rev. Physiol. 61, 243–282 (1999).

49. Moya, A. et al. Rapid acclimation of juvenile corals to CO2-mediated acidification by upregulation of heat shock protein and Bcl-2 genes. Mol. Ecol. 24, 438–452 (2015).

50. Cummings, V. et al. Ocean Acidification at High Latitudes: Potential Effects on Functioning of the Antarctic Bivalve Laternula elliptica. PLOS ONE 6, e16069 (2011).

51. Huan, P., Liu, G., Wang, H. & Liu, B. Identification of a tyrosinase gene potentially involved in early larval shell biogenesis of the Pacific oyster Crassostrea gigas. Dev. Genes Evol. 223, 389–394 (2013).

52. Yang, B. et al. Functional analysis of a tyrosinase gene involved in early larval shell biogenesis in Crassostrea angulata and its response to ocean acidification. Comp. Biochem. Physiol. B Biochem. Mol. Biol. 206, 8–15 (2017).

53. Hüning, A. K. et al. Impacts of seawater acidification on mantle gene expression patterns of the Baltic Sea blue mussel: implications for shell formation and energy metabolism. Mar. Biol. 160, 1845–1861 (2012).

54. Lewontin, R. C. & Hubby, J. L. A Molecular Approach to the Study of Genic Heterozygosity in Natural Populations. II. Amount of Variation and Degree of Heterozygosity in Natural Populations of DROSOPHILA PSEUDOOBSCURA. Genetics 54, 595–609 (1966).

55. Kapsenberg, L. et al. Advancing Ocean Acidification Biology Using Durafet® pH Electrodes. Front. Mar. Sci. 4, (2017).

56. Dickson, A. G., Sabine, C. L. & Christian, J. R. Guide to Best Practices for Ocean CO2 Measurements. (PICES Special Publication, 2007).

57. Gattuso, J.-P., Epitalon, J.-M. & Lavigne, H. seacarb: Seawater Carbonate Chemistry. R package version 3.2.2.

58. Lueker, T. J., Dickson, A. G. & Keeling, C. D. Ocean pCO2 calculated from dissolved inorganic carbon, alkalinity, and equations for K1 and K2: validation based on laboratory measurements of CO2 in gas and seawater at equilibrium. Mar. Chem. 70, 105–119 (2000).

59. Perez, F. F. & Fraga, F. The pH measurements in seawater on the NBS scale. Mar. Chem. 21, 315–327 (1987).

60. Dickson, A. G. Standard potential of the reaction: AgCl(s) + 12H2(g) = Ag(s) + HCl(aq), and and the standard acidity constant of the ion HSO4− in synthetic sea water from 273.15 to 318.15 K. J. Chem. Thermodyn. 22, 113–127 (1990).

61. R Core Team. R: a language and environment for statistical computing.

62. Cosart, T. et al. Exome-wide DNA capture and next generation sequencing in domestic and wild species. BMC Genomics 12, 347 (2011).

63. Bolger, A. M., Lohse, M. & Usadel, B. Trimmomatic: a flexible trimmer for Illumina sequence data. Bioinforma. Oxf. Engl. 30, 2114–2120 (2014).

64. Langmead, B. & Salzberg, S. L. Fast gapped-read alignment with Bowtie 2. Nat. Methods 9, 357–359 (2012).

65. Van der Auwera, G. A. et al. From FastQ data to high confidence variant calls: the Genome Analysis Toolkit best practices pipeline. Curr. Protoc. Bioinforma. 43, 11.10.1–33 (2013).

66. Danecek, P. et al. The variant call format and VCFtools. Bioinformatics 27, 2156–2158 (2011).

67. Kofler, R. et al. PoPoolation: A Toolbox for Population Genetic Analysis of Next Generation Sequencing Data from Pooled Individuals. PLOS ONE 6, e15925 (2011).

68. Storey, J. D., Bass, A. J., Dabney, A. & Robinson, D. qvalue: Q-value estimation for false discovery rate control. (2019).

69. Hivert, V., Leblois, R., Petit, E. J., Gautier, M. & Vitalis, R. Measuring Genetic Differentiation from Pool-seq Data. Genetics 210, 315–330 (2018).

